# Betacoronaviruses Differentially Activate the Integrated Stress Response to Optimize Viral Replication in Lung Derived Cell Lines

**DOI:** 10.1101/2024.09.25.614975

**Authors:** David M. Renner, Nicholas A. Parenti, Susan R. Weiss

## Abstract

The betacoronavirus genus contains five of the seven human viruses, making it a particularly critical area of research to prepare for future viral emergence. We utilized three human betacoronaviruses, one from each subgenus-HCoV-OC43 (embecovirus), SARS-CoV-2 (sarbecovirus) and MERS-CoV (merbecovirus)- to study betacoronavirus interaction with the PKR-like ER kinase (PERK) pathway of the integrated stress response (ISR)/unfolded protein response (UPR). The PERK pathway becomes activated by an abundance of unfolded proteins within the endoplasmic reticulum (ER), leading to phosphorylation of eIF2α and translational attenuation in lung derived cell lines. We demonstrate that MERS-CoV, HCoV-OC43, and SARS-CoV-2 all activate PERK and induce responses downstream of p-eIF2α, while only SARS-CoV-2 induces detectable p-eIF2α during infection. Using a small molecule inhibitor of eIF2α dephosphorylation, we provide evidence that MERS-CoV and HCoV-OC43 maximize replication through p-eIF2α dephosphorylation. Interestingly, genetic ablation of GADD34 expression, an inducible phosphatase 1 (PP1)-interacting partner targeting eIF2α for dephosphorylation, did not significantly alter HCoV-OC43 or SARS-CoV-2 replication, while siRNA knockdown of the constitutive PP1 partner, CReP, dramatically reduced HCoV-OC43 replication. Combining growth arrest and DNA damage-inducible protein (GADD34) knockout with peripheral ER membrane–targeted protein (CReP) knockdown had the maximum impact on HCoV-OC43 replication, while SARS-CoV-2 replication was unaffected. Overall, we conclude that eIF2α dephosphorylation is critical for efficient protein production and replication during MERS-CoV and HCoV-OC43 infection. SARS-CoV-2, however, appears to be insensitive to p-eIF2α and, during infection, may even downregulate dephosphorylation to limit host translation.

**IMPORTANCE:** Lethal human betacoronaviruses have emerged three times in the last two decades, causing two epidemics and a pandemic. Here, we demonstrate differences in how these viruses interact with cellular translational control mechanisms. Utilizing inhibitory compounds and genetic ablation, we demonstrate that MERS-CoV and HCoV-OC43 benefit from keeping p-eIF2α levels low to maintain high rates of virus translation while SARS-CoV-2 tolerates high levels of p-eIF2α. We utilized a PP1:GADD34/CReP inhibitor, GADD34 KO cells, and CReP-targeting siRNA to investigate the therapeutic potential of these pathways. While ineffective for SARS-CoV-2, we found that HCoV-OC43 seems to primarily utilize CReP to limit p-eIF2a accumulation. This work highlights the need to consider differences amongst these viruses, which may inform the development of host-directed pan-coronavirus therapeutics.

## INTRODUCTION

Protein production is critical for cellular survival and viral replication. Translational control offers the cell the chance to respond to various forms of stress that may influence proteostasis or protein quality control. These insults include amino acid starvation, ribosome stalling or collisions, oxidative stress, endoplasmic reticulum (ER) stress, and viral infection. Mammals have evolved an elegant system, termed the integrated stress response (ISR), for detecting and responding to these perturbations and limiting translation while attempting to restore homeostasis (1).

The ISR is a system of four kinases that all converge on the phosphorylation of serine 51 of the alpha subunit of eukaryotic initiation factor 2 (eIF2α). These proteins share highly conserved kinase domains but detect and respond to different types of cellular stress. General control nonderepressible 2 (GCN2), the most ancient ISR kinase conserved down to budding yeast, responds to amino acid starvation, ribosome stalling (1), and ribosome collisions (2). Heme-regulated eIF2α kinase (HRI) senses and responds to heme starvation, oxidative stress (1), and has recently been tied to mitochondrial stress (3). Protein kinase R (PKR) binds to double-stranded RNA (dsRNA), a replication intermediate of RNA and some DNA viruses, making the ISR partly overlap with innate immunity and the interferon response (1, 4). The fourth kinase, PKR-like ER kinase (PERK), is a transmembrane protein residing in the ER. The luminal domain of PERK is bound by binding immunoglobulin proteins (BiP), a chaperone within the ER lumen. As a consequence of ER stress, BiP dissociates from PERK, inducing PERK activation and phosphorylation of eIF2α, which limits translation and the influx of nascent peptides into the ER. PERK, along with inositol requiring enzyme 1α (IRE1α) and activating transcription factor 6 (ATF6), also constitutes part of the unfolded protein response (UPR), which serves to sense and respond to stress within the ER (5). Thus, the ISR serves a central role in detecting and responding to stress within mammalian cells and overlaps extensively with other, more specific stress pathways.

Phosphorylation of eIF2α limits the availability of the eIF2:GTP:Met-tRNA_i_^Met^ ternary complex, thus limiting cap-dependent translation (1, 6). While the translation of most mRNAs is limited when eIF2α is phosphorylated, a subset of mRNAs is translated more efficiently under these conditions. Certain response factors, such as activating transcription 4 (ATF4), have upstream open reading frames (uORFs) in the 5’ end of their mRNAs. During homeostatic conditions, ribosomes preferentially initiate on these uORFs, synthesizing short, abortive peptides rather than the true coding sequence. When ternary complex abundance is low, translation initiation is slowed allowing ribosomes to scan through uORFs or reinitiate on the correct ORF (7). ATF4 is translated under conditions of translation attenuation and serves as the master transcriptional regulator of the ISR. ATF4 induces a transcriptional cascade aimed at alleviating stress and restoring proteostasis. If the stress is too great or cannot be resolved, the ISR can also induce pro-apoptotic genes such as the C/EBP Homologous Protein (CHOP) to destroy chronically stressed cells (8, 9).

If the stress has been resolved, eIF2α must be dephosphorylated to restore full translational capacity. Dephosphorylation is catalyzed by protein phosphatase 1 (PP1), which is directed to p-eIF2α by two different regulatory subunits (10). Constitutive repressor of eIF2α phosphorylation (CReP) directs continuous, low-level dephosphorylation of eIF2α under all conditions (11). This protein serves the role of maintaining a minimal concentration of ternary complex within the cell at all times so that low levels of translation are maintained to respond to stress (1). Growth arrest and DNA-damage inducible 34 (GADD34) is an inducible, uORF-regulated PP1 interacting partner that is induced downstream of ATF4 and highly expressed with prolonged eIF2α phosphorylation (12). This serves as a negative feedback loop within the ISR, promoting robust eIF2α dephosphorylation to restore translation and inhibit GADD34’s own induction if proteostasis has been restored (13) (**Figure 1**).

**Figure 1:**
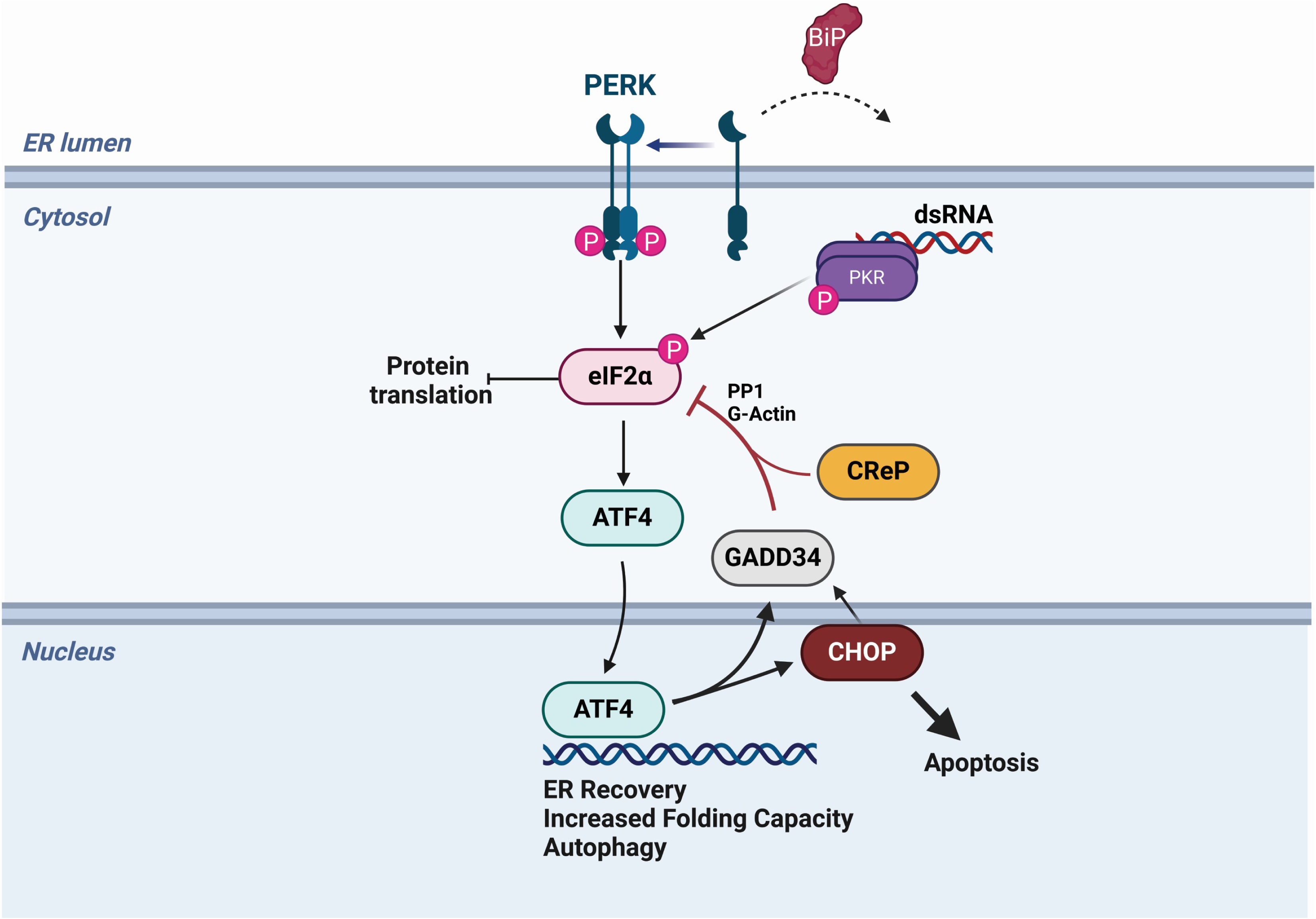
Diagram of the PERK pathway and PKR from the integrated stress response. Following activation of either PERK or PKR, serine 51 on eIF2α is phosphorylated, leading to translational attenuation and the upregulation of ATF4 translation. ATF4 induces a number of recovery responses. GADD34 and CReP promote eIF2α dephosphorylation to restart translation, and CHOP is a a pro-apoptotic transcription factor that promotes death in terminally stressed cells. Created with BioRender.com.

One function of the ISR is to detect and combat viral infection, which has the potential to activate multiple ISR kinases depending on the viral replication cycle. Coronaviruses (CoVs) are large, single-stranded, positive-sense RNA viruses that establish infection within the host ER. To date, there are seven known human CoVs spanning two genera: alpha- and betacoronavirus. In the 21^st^ century, three highly lethal human CoVs have emerged: severe acute respiratory syndrome (SARS)-CoV in 2002, Middle East respiratory syndrome (MERS)-CoV in 2012, and SARS-CoV-2 in 2019. All of these viruses belong to the betacoronavirus genus, but to different subgenera. SARS-CoV and SARS-CoV-2 are sarbecoviruses, while MERS-CoV is a merbecovirus. Furthermore, two common cold causing human coronaviruses – HCoV-OC43 and HCoV-HKU1 – fall into a third subgenus, embecoviruses (14). During infection, CoVs vastly remodel the host ER, form viral replication factories in ER-derived double-membrane vesicles (DMVs) (15-17), and produce dsRNA as a replication intermediate (18). Additionally, three viral structural glycoproteins (spike, membrane, and envelope) are membrane-embedded and require trafficking through the ER, causing the ER to be flooded with viral proteins. Lastly, new viral particles form by budding into the ER-Golgi intermediate complex (ERGIC), thus depleting cellular membranes as new enveloped virions bud from the cell by exocytosis (14). Thus, we hypothesized that coronavirus infection triggers the necessary stress stimuli to induce PKR and PERK activation during infection.

Viral interactions with the ISR have been extensively reported, particularly interactions with PKR. We have previously demonstrated that during infection, MERS-CoV and SARS-CoV-2 interact differently with PKR. MERS-CoV encodes efficient antagonists of PKR activation (19, 20) while SARS-CoV-2 induces p-PKR and p-eIF2α during infection (18). Indeed, many viruses encode antagonists of PKR to limit translational shutdown during infection (21-25), while others have been reported to activate multiple kinases within the ISR (18, 26, 27). Some viruses, such as the alphacoronavirus transmissible gastroenteritis virus (TGEV) (28), herpes simplex 1 (HSV-1) (29), and African swine fever virus (ASFV) (30) even encode GADD34-analogous viral proteins that maintain translation within the infected cell. However, coronavirus interactions with other ISR kinases, such as PERK, have remained relatively unexplored.

Here, we compared three human betacoronaviruses from different subgenera –HCoV-OC43, SARS-CoV-2, and MERS-CoV (31) – and their interactions with the ISR. We focused specifically on the activation of the ISR kinases PERK and PKR, the downstream effects on p-eIF2a, and the role of the eIF2a phosphatases GADD34 and CReP during infection. We found that all three viruses activate PERK during infection, but only SARS-CoV-2 induces p-eIF2α. Despite this, all of these viruses induce downstream signaling events of the ISR, including GADD34 upregulation. Utilizing chemical inhibitors of GADD34 and CReP (32), along with genetic ablation, we show that HCoV-OC43 relies primarily on CReP to maintain eIF2α dephosphorylation and efficient viral replication (1). Disruption of eIF2α dephosphorylation is detrimental to MERS-CoV and, to a greater extent, HCoV-OC43 protein production and replication, but not SARS-CoV-2. Interestingly, our data suggest that SARS-CoV-2 may slow eIF2α dephosphorylation by limiting CReP and GADD34 expression. Our findings elucidate the role of the ISR and p-eIF2α in controlling different human coronavirus infections and establish PP1-mediated eIF2α dephosphorylation (33) as a host-directed therapeutic target for some human betacoronaviruses.

## RESULTS

### HCoV-OC43, SARS-CoV-2, and MERS-CoV activate the unfolded protein response

To understand how different betacoronaviruses interact with the host, we analyzed transcriptomic RNA sequencing (RNA-seq) data from infected A549 lung cell lines expressing either dipeptidyl peptidase 4 (A549^DPP4^) for MERS-CoV or angiotensin converting enzyme 2 (A549^ACE2^) for SARS-CoV-2 and HCoV-OC43 (34). In all infections, we observed a distinct upregulation of the unfolded protein response (UPR) genes, especially PERK-regulated genes. Volcano plots were generated from RNA-seq data from each infection, with select UPR-regulated genes for MERS-CoV (**Figure 2A**), SARS-CoV-2 (**Figure 2B**), and HCoV-OC43 (**Figure 2C**) highlighted in red. HCoV-OC43 infection significantly promoted upregulation of the largest number of UPR-related genes (**Figure 2C**) compared to SARS-CoV-2 (**Figure 2B**) or MERS-CoV (**Figure 2A**). However, all three viruses strongly upregulated three UPR genes (labeled in **Figure 2**), the PERK/ISR-regulated genes *ATF3* (35); DNA-damage inducible transcription factor 3 (*DDIT3)*, encoding CHOP; and *PPP1R15A, encoding GADD34* (1). As we recently reported, SARS-CoV-2 failed to induce IRE1α-regulated genes (**Figure 2B**) while MERS-CoV and HCoV-OC43 did (34). Gene set enrichment analysis (GSEA) also showed significant upregulation of UPR-related genes during MERS-CoV (**Figure 2D**) and HCoV-OC43 (**Figure 2F**) infection, while SARS-CoV-2 (**Figure 2E**) displayed non-significant enrichment.

**Figure 2:**
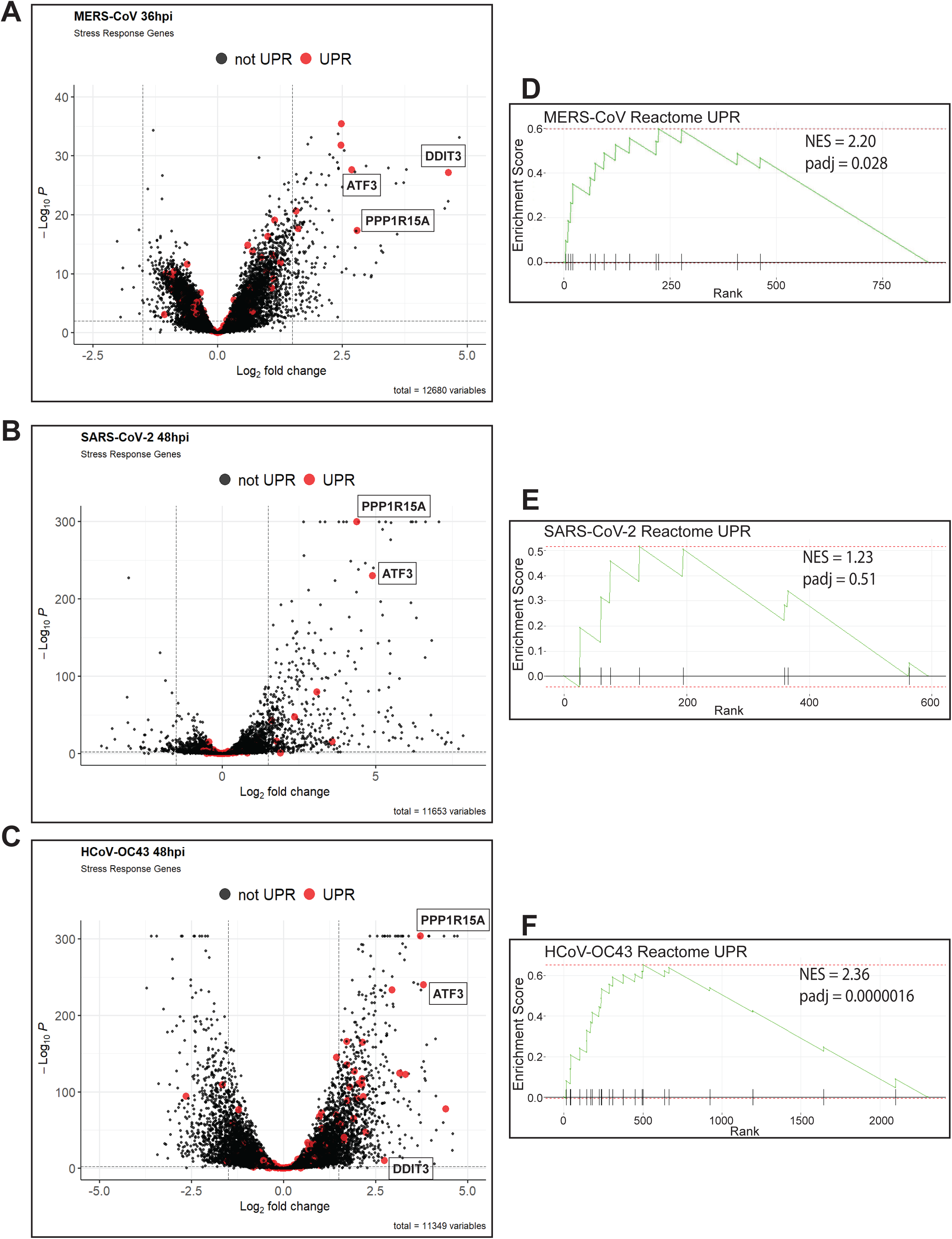
MERS-CoV, SARS-CoV-2, and HCoV-OC43 display signature of PERK and UPR activation. (A-C) RNA-seq datasets of MERS-CoV infection in A549DPP4 cells at 36hpi (A), SARS-CoV-2 (B) or HCoV-OC43 (C) infection in ACE2-A549 cells at 48hpi were compared to mock infections and differentially expressed genes called using DESeq2. UPR-regulated gene highlighted (in red) volcano plots were generated using EnhancedVolcano. (D-F) Gene Set Enrichment Analysis (GSEA) using the RNA-seq datasets for B-D. Pathway enrichment plots for the Reactome Unfolded Protein Response (UPR) gene list were generated for MERS-CoV (D), SARS-CoV-2 (E), and HCoV-OC43 (F) infected A549s. Normalized enrichment score (NES) and p-adjusted value (padj) are displayed on the plots.

### MERS-CoV and HCoV-OC43 do not induce p-eIF2α despite PERK activation

To confirm that PERK is activated during infection by these betacoronaviruses, A549 cells expressing the appropriate viral receptor were infected at a multiplicity of infection (MOI) of 5. In addition to SARS-cov-2, OC43 and MERS-coV we also included MERS-CoV-nsp15^mut^/ΔNS4a, an immunostimulatory double mutant encoding a catalytically inactive endoribonuclease in the nsp15 protein and a deletion of the NS4a encoded protein (nsp15^mut^/ΔNS4a) (20). Whole cell lysates were collected at 24, 48, and 72 hours post-infection (hpi) for immunoblot analysis. Due to the lack of effective phospho-PERK antibodies for human samples, PERK activation was assessed using Phos-tag^TM^ SDS-PAGE, which slows the migration of phosphorylated proteins through the polyacrylamide, thus separating phosphorylated and unphosphorylated species. As positive controls, cells were treated with thapsigargin (Tg), a SarcoEndoplasmic Reticulum Calcium ATPase (SERCA) inhibitor (36), for one hour or tunicamycin (TM), an N-linked glycosylation inhibitor (34), for eight hours to induce ER stress. These conditions showed clear separation between phosphorylated and non-phosphorylated PERK bands (**Figure 3A-C**). Lysates from cell infected with all the viruses examined showed an upper band in these blots representing p-PERK, demonstrating PERK activation during infection. PERK activation can also be visualized by standard SDS-PAGE, with virus-infected cells or cells treated with either Tg or TM. A band shift and shading pattern is observed indicating PERK phosphorylation and activation (**Figure 3D-F**). This led us to conclude that all three viruses activate PERK during infection.

**Figure 3:**
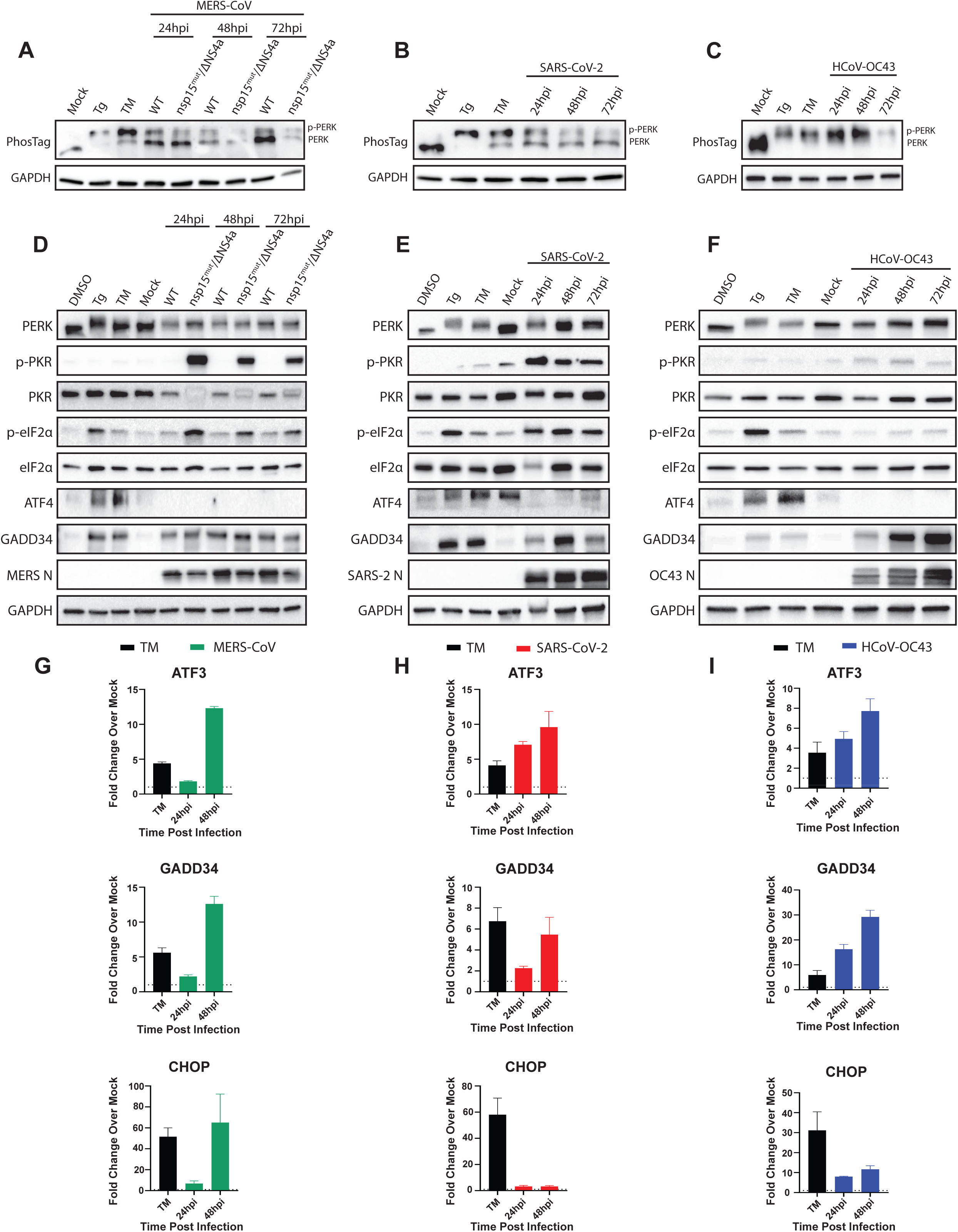
MERS-CoV, SARS-CoV-2, and HCoV-OC43 all activate PERK and downstream signaling during infection. (A-F). In all blots, thapsigargin (Tg,1μM treatment for 1 hour) and tunicamycin (TM, 1μg/mL treatment for 8 hours) served as positive controls, while DMSO (0.1%) served as a vehicle control. Cells (A - A549^DPP4^, B and C A549^ACE2^) were infected with the indicated viruses or mock infected and whole-cell lysates collected at the indicated timepoints. (A-C) Extracted proteins were resolved in SDS-polyacrylamide gels containing 50μM Phosbind acrylamide and Mn2+ to separate phosphorylated and unphosphorylated proteins. Gels were transferred and immunoblotted for PERK (top gel – PhosTag). GAPDH run by standard SDS-PAGE served as a loading control. (D-F) Western immunoblots were performed by standard SDS-PAGE for the indicated proteins. (G-I) Cells were treated with DMSO or 1μg/mL tunicamycin (TM) for 8 hours before total RNA was extracted. (G) A549DPP4 cells were mock infected or infected at MOI 5 with MERS-CoV and total RNA extracted at the indicated timepoints. (H and I) A549ACE2 cells were mock infected or infected with SARS-CoV-2 (H) or HCoV-OC43 (I) at MOI 5 and total RNA collected at the indicated timepoints. Expression of the indicated genes was determined using RT-qPCR, with fold change over mock values being calculated as 2-Δ(ΔCt).

As we previously reported, wild type (WT) MERS-CoV fails to induce PKR activation (indicated by PKR phosphorylation) or eIF2α phosphorylation up to 72hpi (**Figure 3D**) (18). MERS-CoV-nsp15^mut^/ΔNS4a strongly induced p-PKR and p-eIF2α throughout the course of infection as we reported previously (20), confirming that parental MERS-CoV effectively antagonizes PKR to limit eIF2α phosphorylation. Similar to WT MERS-CoV, HCoV-OC43 (**Figure 3F**) also failed to activate PKR or induce p-eIF2α during infection, although the mechanism of PKR antagonism remains unclear. However, SARS-CoV-2 robustly activated PKR and induced p-eIF2α over the course of infection (**Figure 3E**) (18).

It is striking that, despite activation of at least one ISR kinase during infection and apparent ISR gene induction, WT MERS-CoV (**Figure 3D**) and HCoV-OC43 (**Figure 3F**) still fail to induce p-eIF2α during infection. To further assess ISR activation we next examined ATF4 expression during infection, which should occur rapidly following eIF2α phosphorylation (7). As expected, ATF4 is readily detectable in cells treated with either thapsigargin or tunicamycin. However, during infection with any of the three viruses, with or without the presence of p-eIF2α, ATF4 could not be detected at any timepoint (**Figures 3D-3F**). This has been reported previously by other groups probing for ATF4 during infections with coronaviruses (37, 38), however, it is still unclear why this occurs. Despite the absence of detectable ATF4 during infection with any virus, ATF4-regulated genes were highly upregulated. MERS-CoV (**Figure 3G**) and HCoV-OC43 (**Figure 3I**) both induced ATF3, GADD34, and CHOP at increasing levels over the course of infection. While HCoV-OC43 induced much higher levels of GADD34 compared to MERS-CoV, CHOP induction by MERS-CoV dwarfed the other viruses, matching recent reports that MERS-CoV strongly induces apoptosis through PERK and CHOP signaling (39, 40). Interestingly, SARS-CoV-2 (**Figure 3H**) also induced ATF3 and GADD34 throughout the course of infection but failed to significantly upregulate CHOP. This indicates that, while PERK activation and signaling is a common feature of betacoronavirus infection, there are differences (maybe more than nuances?) in the induction of certain responses that remain to be explored.

To understand the absence of eIF2α phosphorylation despite PERK activation during MERS-CoV and HCoV-OC43 infection, we probed for GADD34 protein expression. GADD34 was translated following Tg or TM treatment, confirming that this pathway can be induced in as little as 1 hour following ER stress. Consistent with the transcriptional induction of GADD34 (**Figure 3G-I**), GADD34 protein expression was also observed over the course of MERS-CoV, SARS-CoV-2, and HCoV-OC43 infection (**Figure 3D-F**). This suggested that GADD34 expression during WT MERS-CoV and HCoV-OC43 infection may be keeping p-eIF2α levels below the limit of detection for immunoblotting. The ability of cells to dephosphorylate eIF2α during TM treatment has been noted in the literature (41), and demonstrates that GADD34 is capable of promoting dephosphorylation of eIF2α despite continued ER stress.

### Betacoronaviruses promote translational shutoff with or without p-eIF2α

To understand the impact on overall translation in cells infected with each betacoronavirus, we utilized puromycin incorporation to visualize nascent peptide production. Cells were infected with each virus at a MOI of 5 and, at the indicated timepoints, puromycin was added to the media for incorporation into nascent peptide chains. Whole-cell lysates were then collected and subjected to immunoblotting stained with an antibody raised against puromycin as a measure of total protein translation and with viral nucleocapsid (N) antibody, which served as a marker of infection and a readout of viral protein synthesis (19). Tg treatment served as a positive control for ER stress and p-eIF2α-mediated translational attenuation.

**Figure 4A** shows infection with WT MERS-CoV or the MERS-CoV nsp15^mut^/ΔNS4a double mutant virus that induces p-eIF2α during infection (20) (see **Figure 3D**). Immunoblots for puromycin incorporation revealed that WT MERS-CoV produces a progressive shutdown of host translation despite the lack of p-eIF2α during infection, while conversely, viral translation of N increased over the course of infection. MERS-CoV-nsp15^mut^/ΔNS4a, which activates PKR and induces p-eIF2α during infection, promotes a faster translational shutoff, supporting a role of p-eIF2α in limiting translation during CoV infection. However, both viruses appear to reach similar levels of translational attenuation at late times post infection. In contrast to the progressive translational shutoff induced by WT MERS-CoV infection, SARS-CoV-2 appears to rapidly reduce host translation to very low levels within 24 hours of infection, with puromycin incorporation remaining low at all timepoints examined (**Figure 4B**). However, SARS-CoV-2 N, similar to MERS-CoV N, continues to be translated despite very low levels of global translation within infected cells. HCoV-OC43 infection also induced a rapid shutoff of translation within infected cells that was similar to the attenuation induced by Tg treatment (**Figure 4C**). This was surprising because HCoV-OC43, like WT MERS-CoV, fails to induce p-eIF2α during infection.

**Figure 4:**
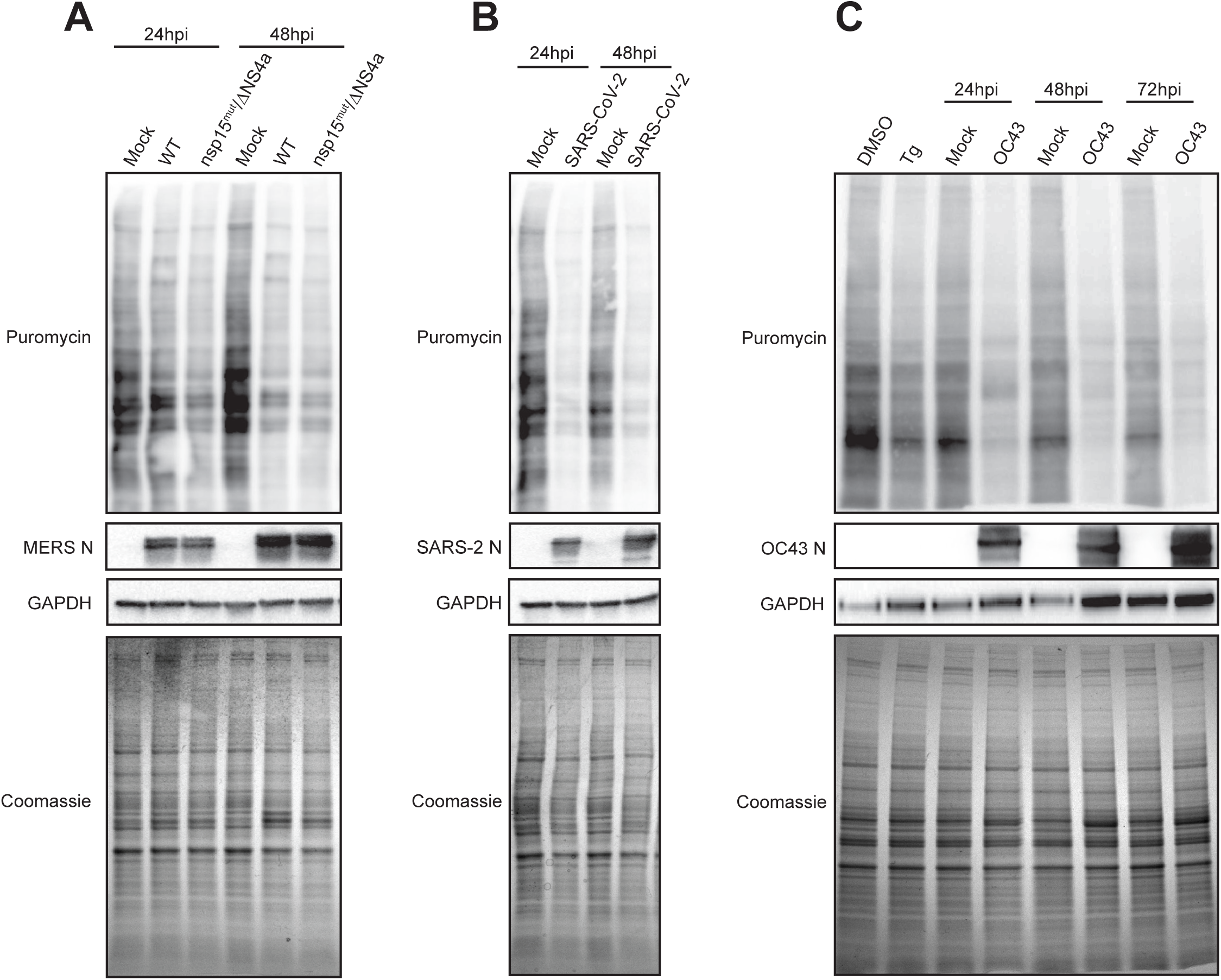
Global translation during betacoronavirus infection. A549 cells expressing the appropriate viral receptors were treated with 0.1% DMSO, 1μM thapsigargin (Tg) for 1 hour, mock infected, or infected at MOI=5. At the indicated times, 10μg/mL of puromycin was added to cells for 10 minutes before lysis and total protein collection. Samples were subjected to immunoblotting for the indicated proteins, while Coomassie staining was used as a readout of total protein. (A) MERS-CoV or MERS-CoV nsp15mut/ΔNS4a infected A549DPP4 cells. (B) SARS-CoV-2 infected A549ACE2 cells. (C) A549ACE2 infected HCoV-OC43 cells. N = Nucleocapsid protein.

### GADD34 Is a Druggable Target During Betacoronavirus Infection

Since WT MERS-CoV and HCoV-OC43 both limit eIF2α phosphorylation during infection, and p-eIF2α is detrimental to MERS-CoV infection (20), we asked if inhibition of GADD34 during betacoronavirus infection would limit viral replication. GADD34 has been reported to be inhibited by several compounds that target the GADD34:PP1 holoenzyme (42). Of these, salubrinal (32) has been utilized widely in the literature. Therefore, salubrinal was used during infection to test its therapeutic potential against betacoronaviruses.

We began by demonstrating that salubrinal is sufficient to induce p-eIF2α during CoV infection. 15μM of salubrinal has been reported as the approximate EC_50_ value for inhibiting the GADD34:PP1 holoenzyme in cells (32), and 20μM has been commonly used in the literature (43, 44) and is the dose utilized in this study. We compared HCoV-OC43 and SARS-CoV-2 because they displayed different eIF2α phenotypes while able to replicate within the same A549^ACE2^ cell line. Cells were mock infected or infected with HCoV-OC43 or SARS-CoV-2 at MOI= 5 PFU/cell and incubated for 24 hours to establish viral infection before salubrinal or Sal003 (43), a salubrinal derivative with similar function, was added for 4 or 24 hours. Whole-cell lysates were collected and analyzed by immunoblot (**Figure 5A-B**). HCoV-OC43 and SARS-CoV-2 activated PERK and induced GADD34 expression with or without inhibitor treatment. However, only salubrinal or Sal003 treatment induced p-eIF2α during infection, confirming that this inhibitor can promote p-eIF2α (**Figure 5A**). Immunoblots of SARS-CoV-2-infected cells demonstrated no difference in p-eIF2α induction, which was present in treated or untreated infections (**Figure 5B**). We next examined the impact on viral protein production following prolonged treatment with 20μM of salubrinal in A549 cells. To do so, we used immunoblotting of viral N, the most abundant viral protein in infected cells, as a readout for viral translation. HCoV-OC43 showed high sensitivity to salubrinal, producing almost no detectable N protein over the course of infection (**Figure 5C**). We also investigated the impact of salubrinal treatment on HCoV-OC43 replication by treating cells infected immediately after infection. HCoV-OC43 titers were reduced by approximately 10-fold at 24hpi with salubrinal treatment and 100-fold at 48hpi and 72hpi (**Figure 5E**). In contrast, SARS-CoV-2 infections demonstrated a steady increase in N levels in A549^ACE2^ cells with or without salubrinal treatment and no defect in viral replication (**Figure 5D** and **E**). Similar treatments in A549^DPP4^ cells infected with MERS-CoV or MERS-CoV nsp15^mut^/ΔNS4a were performed (**Figure S1**). Without salubrinal treatment, we observed a steady increase of N abundance over the course of infection with both viruses, indicating efficient translation. However, treatment of infected cells with salubrinal drastically reduced N abundance during WT MERS-CoV infection. The levels of N produced by MERS-CoV-nsp15^mut^/ΔNS4a were reduced by salubrinal treatment to an even greater extent than WT virus. Examining MERS-CoV replication, MERS-CoV-nsp15^mut^/ΔNS4a is attenuated (20), displaying 2 to 5-fold fewer PFU/mL released at each timepoint compared to WT virus (**Figure S1A**). Salubrinal treatment reduced WT MERS-CoV and MERS-CoV nsp15^mut^/ΔNS4a titers by 10 to 100-fold at each timepoint examined in A549^DPP4^ cells. Overall, these suggdata demonstrate that replication of MERS-CoV and HCoV-OC43 is sensitive to salubrinal treatment and inhibition of eIF2α dephosphorylation during infection, while SARS-CoV-2 is not.

**Figure 5:**
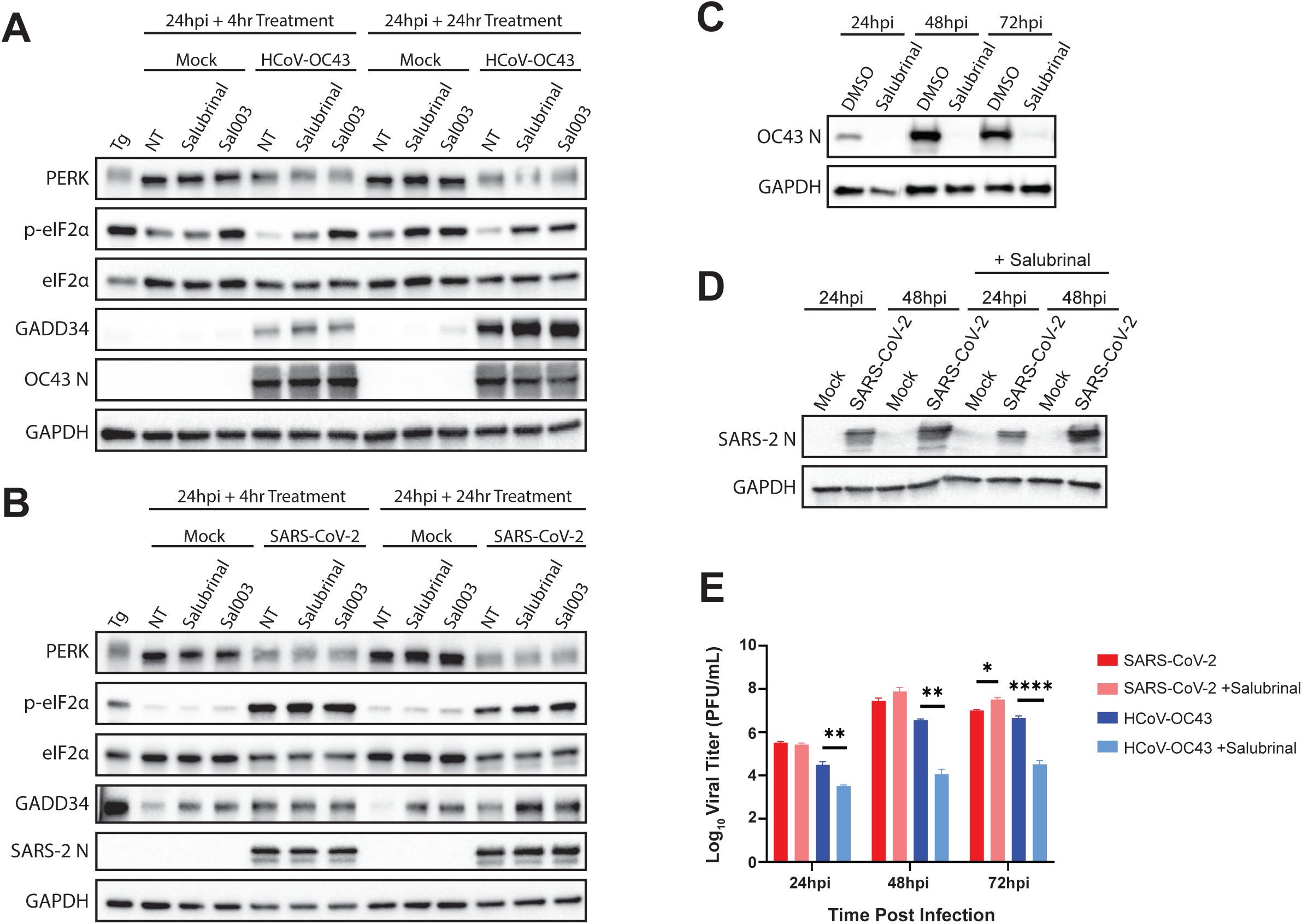
Salubrinal treatment is effective against HCoV-OC43, but not SARS-CoV-2. A549^ACE2^ cells were mock infected or infected at MOI=5 with HCoV-OC43 (A, C, E) or SARS-CoV-2 (B, D, E). (A and B) At 24hpi, cell media was replaced with media containing 20μM salubrinal or 20μM Sal003, and infections were allowed to progress for 4 or 24 more hours. At the indicated timepoints, whole-cell lysates were collected. Immunoblotting was performed for the indicated proteins. NT = no treatment. Thapsigargin (Tg, 1μM for 1 hour) was used as a positive control for p-eIF2α. (C-E) A549^ACE2^ cells were mock infected or infected with the indicated viruses at MOI=5 and treated immediately after virus absorption with 20μM salubrinal or 0.1% DMSO. At the indicated timepoints, cells were lysed and whole-cell lysates collected. Immunoblotting was performed to probe for the indicated proteins, with viral N serving as a readout of viral translation. HCoV-OC43 blots (C) and SARS-CoV-2 blots (D) are shown. (E) A549^ACE2^ cells were infected with HCoV-OC43 or SARS-CoV-2 at MOI = 0.1. Immediately following virus absorption, cells were treated with 20μM salubrinal or 0.1% DMSO. At the indicated timepoints, supernatants were collected and infectious virus quantified by plaque assay. Statistics were calculated using 2-way ANOVA. * = p < 0.05; ** = p < 0.01; *** = p < 0.001; **** = p < 0.0001; ns = not significant.

### GADD34 knockout only slightly impacts HCoV-OC43 replication

To validate our results using salubrinal, we utilized CRISPR-Cas9 in our A549^ACE2^ cells to knock out GADD34 or introduced a control, scrambled single guide RNA (sgRNA). *GADD34 knockout (KO) was validated using genomic DNA sequencing*, GADD34 expression by immunoblot and by testing translational recovery during Tg treatment (**Figure S2**). As seen in **Figure S2A**, control sgCtrl cells produce GADD34 protein and begin to recover translation after only two hours of Tg treatment, with levels of translation steadily increasing over four hours. In contrast, GADD34 KO cells fail to produce GADD34 protein or restart translation at any point (**Figure S2B**), confirming the loss of GADD34. Two separate GADD34 KO clones (clone 15 and clone 23) were chosen for infection with either HCoV-OC43 or SARS-CoV-2.

The sgCtrl generated clone and both GADD34 KO clones were infected with SARS-CoV-2 or HCoV-OC43 at a MOI of 5. Infected cells showed robust PERK activation, as assessed by immunoblot analysis of whole-cell lysates harvested from cells following treatment with Tg or infection with HCoV-OC43 (**Figure 6A**) or SARS-CoV-2 (**Figure 6C**). Phosphorylation of eIF2α was also detected following Tg treatment and over the course of SARS-CoV-2 infection (**Figure 6C**). No consistent impact on SARS-CoV-2 infectious virus production was observed (**Figure 6D**). However, p-eIF2α was not induced during HCoV-OC43 infection of sgCtrl cells nor in infections of both GADD34 KO clones (**Figure 6A**). Thus, GADD34 KO does not appear to significantly alter the phosphorylation state of eIF2α during HCoV-OC43 or SARS-CoV-2 infection. Loss of GADD34 also failed to consistently reduce HCoV-OC43 titers in either knockout clone, suggesting that our hypothesis regarding the role of GADD34 in HCoV-OC43 infection to be incorrect (**Figure 6B**).

**Figure 6:**
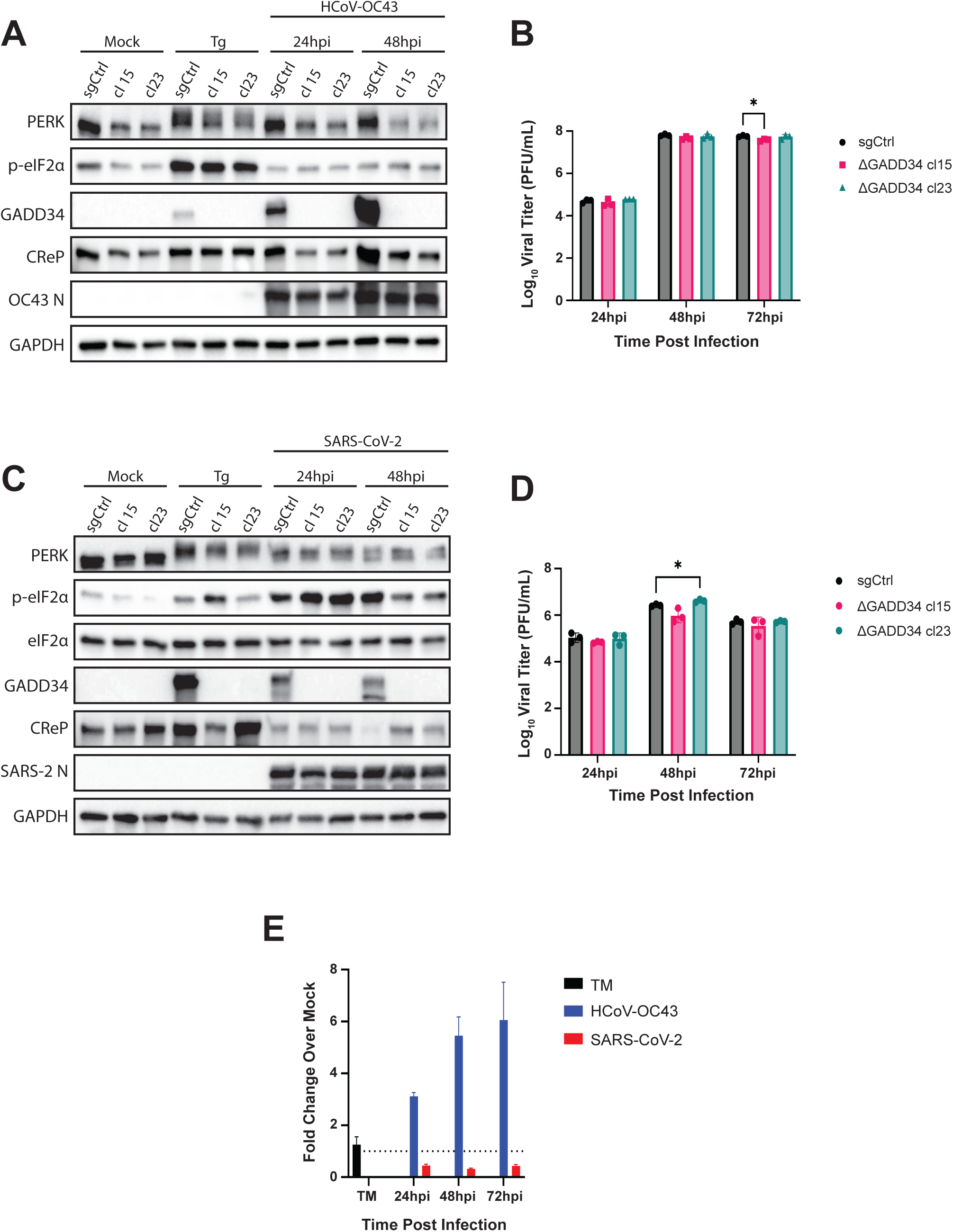
GADD34 knockout has little impact on HCoV-OC43 and SARS-CoV-2 replication. (A-F) Single cell clones of A549^ACE2^ cells with either a nontargeting (sgCtrl) or GADD34 targeted sgRNAs were mock infected or infected at MOI = 2 with HCoV-OC43 (A and B) or SARS-CoV-2 (C and D). At the indicated timepoints, whole-cell lysates (A and C) or RNA (E) was collected. (A and B) Western immunoblot for the indicated proteins. Thapsigargin (Tg, 1h at 1μM) was used as a positive control for ER stress. (A) HCoV-OC43 infections. (C) SARS-CoV-2 infections. (B and D) At the indicated timepoints, supernatants from infected cells were collected and infectious virus quantified by plaque assay. (B) HCoV-OC43 infections. (D) SARS-CoV-2 infections. (E) Total RNA was used for RT-qPCR of CReP transcripts. Values are displayed as fold change over mock, calculated by 2^-Δ(ΔCt)^. Statistics by 2-way ANOVA. * = p < 0.05.

It is surprising that GADD34 KO is not as effective as salubrinal, a known GADD34 inhibitor, at reducing HCoV-OC43 titers. While salubrinal has been reported to inhibit GADD34 (32, 45, 46), it has also been reported to inhibit the PP1 holoenzyme in complex with CReP (32). Thus, the additional efficacy of salubrinal may be due to co-inhibition of CReP during HCoV-OC43 infection. We investigated CReP expression at the RNA level by RT-qPCR and at the protein level by immunoblotting of lysates from cells infected with HCoV-OC43 or SARS-CoV-2. Surprisingly, we observed a dramatic increase in CReP mRNA levels during HCoV-OC43 infection (3-8-fold) (**Figure 6E**) as well as a dramatic increase in CReP protein levels (**Figure 6A**). Conversely, SARS-CoV-2 promoted reduced CReP expression at the RNA and protein expression levels during infection (**Figures 6E** and **6D**).

### CReP Is necessary for efficient HCoV-OC43 replication

To investigate the role of CReP in betacoronavirus replication, we utilized siRNA to knockdown (KD) CReP expression in A549^ACE2^ cells before infecting with HCoV-OC43 or SARS-CoV-2. CReP protein levels were efficiently reduced compared to treatment with a scrambled siRNA control (siCtrl) (**Figure 7A** and **7C**). 72 hours after siRNA transfection, cells were infected with HCoV-OC43 or SARS-CoV-2 for the indicated times and whole-cell lysates collected for immunoblots. During infection of siCtrl-treated cells with HCoV-OC43, we observed a decrease in p-eIF2α levels below background of mock infected siCtrl cells. Knockdown of CReP during infection was maintained through the course of infection and led to an increase in p-eIF2α levels, particularly at 24hpi (**Figure 7A**). This increase in p-eIF2α at 24hpi also corresponded with a notable decrease in HCoV-OC43 N protein. CReP KD also reduced HCoV-OC43 titers by approximately 100-fold at 24hpi, with the defect decreasing to only 10-fold at 48hpi and 3-fold at 72hpi (**Figure 7B**). We hypothesize that this diminishing effect viral replication as the infection progresses may be due to CreP upregulation paired with siRNA turnover. These data, as well as the significant impact of salubrinal treatment on HCoV-OC43 replication (**Figure 5E**), leads us to conclude that HCoV-OC43 preferentially promotes eIF2α dephosphorylation and viral replication through CreP rather than GADD34.

**Figure 7:**
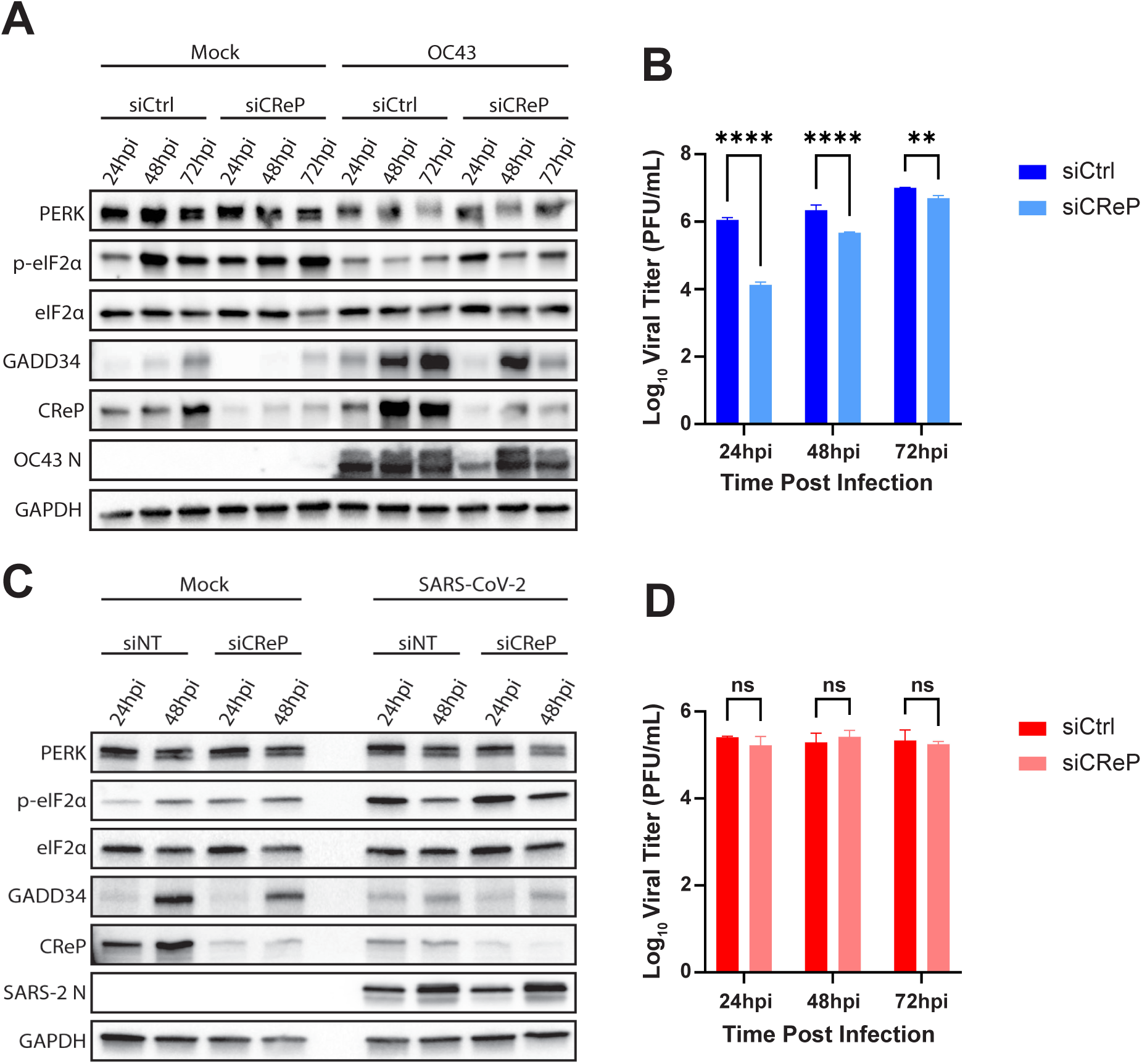
CReP knockdown reduces HCoV-OC43, but not SARS-CoV-2, replication. A549^ACE2^ cells were treated with siRNA targeting CReP (siCReP) or a scrambled control (siCtrl) for 72 hours before mock infection or infection with HCoV-OC43 (A and B) or SARS-CoV-2 (C and D) at an MOI = 2. At the indicated timepoints, whole-cell lysates (A and C) or supernatants from infected cells (B and D) were collected. (A and C) Western immunoblots for the indicated proteins from HCoV-OC43 infected cells (A) or SARS-CoV-2 infected cells (C). (B and D) Viral titers were quantified by plaque assay for HCoV-OC43 (B) or SARS-CoV-2 (D) in the indicated conditions. Statistics by 2-way ANOVA. * = p < 0.05; ** = p < 0.01; *** p < 0.001; **** = p < 0.0001.

In contrast to OC43 infection, CreP KD during SARS-CoV-2 infection failed to have a major impact on p-eIF2α levels. Due to cell death at the MOI used, a small decrease in p-eIF2α levels at 48hpi with both CreP KD and siCrtl was observed. (**Figure 7C**). This KD of CreP failed to reduce SARS-CoV-2 replication (**Figure 7D**), supporting the ability of SARS-CoV-2 to circumvent cellular translational control.

### CreP and GADD34 both contribute to HCoV-OC43 replication

Having examined the individual contributions of GADD34 and CreP to betacoronavirus replication, we next combined these conditions to determine if CreP KD and GADD34 KO would have a combinatorial effect on HCoV-OC43 replication. To do this, we treated sgCtrl A549^ACE2^ cells or GADD34 KO cells (clone 23) with scrambled (siCtrl) or CReP targeting (siCReP) siRNA. Immunoblots of whole-cell lysates harvested from HCoV-OC43-infected cells at 24hpi (**Figure 8A**) and 48hpi (**Figure 8B**) were performed. As expected, GADD34 expression was ablated in GADD34 KO cells while CReP expression was efficiently reduced with siRNA treatment in either cell line at both timepoints. As observed in **Figure 7**, CReP KD in either sgCtrl or GADD34 KO cells led to an increase in p-eIF2α during HCoV-OC43 infection at 24hpi and 48hpi (**Figure 8A** and **8B**). Additionally, GADD34 KO alone did not lead to increased p-eIF2α phosphorylation levels (**Figure 8A** and **8B**) and failed to impact HCoV-OC43 replication (**Figure 8E**). In contrast, CReP KD in sgCtrl cells significantly reduced HCoV-OC43 titers by nearly 50-fold at 24hpi, with this difference again diminishing at later timepoints. However, combining CReP KD in GADD34 KO cells led to an even greater decrease in HCoV-OC43 titers, reducing viral replication another 5-fold compared to CReP KD alone (**Figure 8E**). This difference was sustained, but again diminished by 48 and 72hpi. Together, these data suggest that while CReP is the main driver for eIF2α dephosphorylation and HCoV-OC43 replication, GADD34 also plays a role in further boosting viral replication.

**Figure 8:**
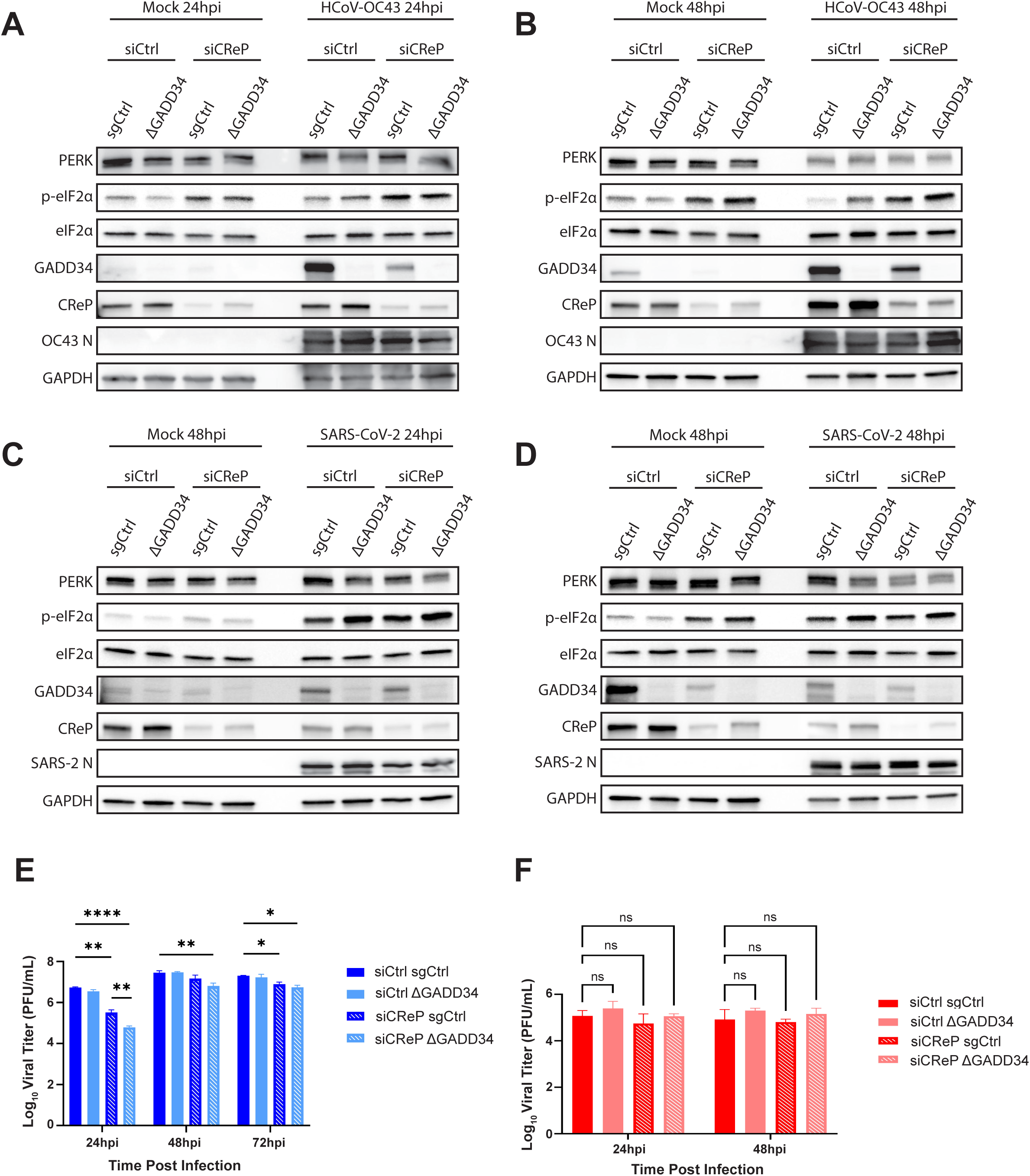
CReP knockdown in GADD34 knockout cells has a combinatorial effect on HCoV-OC43 replication. A549^ACE2^ sgCtrl cells or GADD34 KO cells (clone 23 – ΔGADD34) were treated with control siRNA (siCtrl) or CReP-targeting siRNA (siCReP) for 72 hours before being infected with HCoV-OC43 (A, B, E) or SARS-CoV-2 (C, D, F). At the indicated timepoints, whole-cell lysates (A-D) or supernatants (E and F) were collected. (A-D) Western immunoblots for the indicated proteins were performed from HCoV-OC43 infected cells (A and B) or SARS-CoV-2 infected cells (C and D). (E and F) Infectious virus was quantified by plaque assay from HCoV-OC43 infected samples (E) and SARS-CoV-2 infected samples (F). Solid lines indicated siCtrl treatment while dashed lines represent siCReP treatment. Statistics by 2-way ANOVA. * = p < 0.05; ** = p < 0.01; *** p < 0.001; **** = p < 0.0001.

As expected, neither CReP KD, GADD34 KO, nor the combination of the two significantly altered the induction of p-eIF2α during SARS-CoV-2 infection at 24hpi (**Figure 8C**) or 48hpi **(Figure 8D**). Additionally, despite the loss of GADD34, the reduction in CReP, or a combination of the two, SARS-CoV-2 replication was again unchanged under any condition tested (**Figure 8F**).

## DISCUSSION

We have presented evidence that HCoV-OC43, SARS-CoV-2, and MERS-CoV – representing different betacoronavirus subgenera (31) – activate the PERK arm of the UPR. In **Figure 2**, we utilized RNA-seq data from infections of A549 cells with each virus (34) to demonstrate enrichment of ISR-regulated genes, including *ATF3* (35), GADD34 (gene name *PPP1R15A*), and CHOP (gene name *DDIT3*) (1). We have previously shown that MERS-CoV effectively antagonizes PKR during infection and fails to induce phosphorylation of eIF2α, while SARS-CoV-2 infection activates PKR and induces p-eIF2α (18). However, we have also shown that cells lacking PKR still phosphorylate eIF2α during SARS-CoV-2 infection suggesting at least one other ISR kinase is active (18). Due to the remodeling of the host ER during coronavirus infection (14) and evidence from other groups that overexpression of spike protein alone is sufficient to induce the UPR (37, 47), we hypothesized that PERK activation during infection with these viruses was contributing to the responses observed in our RNA-seq data.

Despite confirming PERK activation and downstream signaling during CoV infection (**Figure 3**), we observed that WT MERS-CoV and HCoV-OC43 failed to induce detectable p-eIF2α during infection. Having shown the induction of GADD34 during infection with each virus at the mRNA and protein levels, the most parsimonious explanation for this disconnect is that GADD34 is driving eIF2α dephosphorylation (13) during WT MERS-CoV and HCoV-OC43 infection. Indeed, our positive controls Tg and TM reveal this process in action in A549 cells. As shown in **Figure 3D-3F**, 1 hour of Tg treatment is sufficient to activate PERK, induce p-eIF2α, and promote ATF4 and GADD34 translation. Eight hours of TM treatment similarly induces PERK activation and ATF4/GADD34 translation. However, at this timepoint there is no longer detectable p-eIF2α because enough GADD34 has accumulated to now dephosphorylate eIF2α. Such instances of viruses preferring the dephosphorylated state of eIF2α have been observed with pseudorabies virus where characterization of viral proteins with similar functions to GADD34 demonstrate the need to maintain translation during infection (48-50). However, we and others have shown that coronaviruses mediate host translational shutdown during infection (**Figure 4**) using non-structural protein (nsp)1 (51-55), even without the induction of p-eIF2α, which is detrimental to MERS-CoV replication and protein production (20). It is thus intriguing that SARS-CoV-2 shows efficient N production despite continuous phosphorylation of eIF2α during infection (**Figure 4B**). This suggests that MERS-CoV and HCoV-OC43, but not SARS-CoV-2, require a specific translational context within the infected cell to replicate optimally.

To test the importance of eIF2α dephosphorylation on betacoronavirus infection, we utilized salubrinal, a widely used inhibitor of eIF2α dephosphorylation. This compound has been reported to target the PP1:GADD34 and PP1:CReP holoenzymes to disrupt eIF2α dephosphorylation (32, 42), thus making it a potential host-directed antiviral for coronavirus infection. We found that salubrinal treatment of A549 cells is effective against HCoV-OC43 (**Figure 5**) and MERS-CoV (**Figure S1**) replication and protein production. However, SARS-CoV-2 showed little if any sensitivity to salubrinal treatment (**Figure 5D-5E**). It is unclear what could be mediating this difference, and much more research will be required to uncover the exact mechanism. It is also interesting that the extreme sensitivity of HCoV-OC43 to salubrinal treatment seems to distinguish this common cold coronavirus from the lethal human coronaviruses.

Due to the nonspecific nature of small molecule inhibitors, we utilized a CRISPR-Cas9 KO of GADD34 to confirm its role in HCoV-OC43 and SARS-CoV-2 infection. Due to the similar phenotypes between HCoV-OC43 and MERS-CoV, and the ability of HCoV-OC43 to infect the same A549^ACE2^ cell line as SARS-CoV-2, we proceeded to compare only HCoV-OC43 and SARS-CoV-2. In contrast to our initial hypothesis, GADD34 KO cells showed no detectable alterations in p-eIF2α levels (**Figure 6A** and **6C**) or viral replication (**Figure 6B** and **6D**) during HCoV-OC43 or SARS-CoV-2 infection. These results are supported by similar findings that were recently published (56), although we have further expanded upon this to provide a potential explanation for our shared negative results. A dramatic increase in CReP mRNA and protein levels was observed during HCoV-OC43 infection (**Figures 6E** and **6A**), while a reduction of both was seen during SARS-CoV-2 infection (**Figures 6E** and **6C**). Thus, our data suggest that CReP, another target of salubrinal (32), is the main driver of eIF2α dephosphorylation during HCoV-OC43 infection.

Supporting the role of CReP in dephosphorylation of eIFα, we found that knocking down CReP expression using siRNA led to increased p-eIF2α levels, decreased N expression (**Figure 7A**), and a significant reduction in viral titers (**Figure 7B**) during HCoV-OC43 infection. SARS-CoV-2 replication (**Figure 7D**) and p-eIF2α levels (**Figure 7C**) once again remained unchanged. To understand if GADD34 and CReP are working cooperatively during HCoV-OC43 infection, a GADD34 KO combined with a CReP KD was performed. These data clearly show a combinatorial role for these PP1 binding partners during HCoV-OC43 infection due to CReP KD in GADD34 KO cells having a more dramatic effect on HCoV-OC43 replication than CReP KD alone (**Figure 8E**). Thus, we conclude that CReP is the primary factor for promoting dephosphorylation of eIF2α during HCoV-OC43 infection, but that GADD34 also plays a role in optimizing HCoV-OC43 replication. In contrast to this, SARS-CoV-2 was still unaffected by the combined loss of GADD34 and CReP (**Figure 8F**), and p-eIF2α levels were unaltered during infection of any condition (**Figures 8C** and **8D**).

We thus conclude that HCoV-OC43 and SARS-CoV-2 have diverged in their reliance on host translational control via eIF2α phosphorylation. HCoV-OC43 appears to employ multiple mechanisms to limit eIF2α phosphorylation, including antagonizing PKR (**Figure 3F**), upregulating GADD34 (**Figure 3I**) and CReP (**Figure 6E**), and promoting eIF2α dephosphorylation (**Figure 8A** and **8B**). SARS-CoV-2, however, diverges from HCoV-OC43 in all of these aspects and promotes sustained eIF2α phosphorylation throughout the course of infection (**Figure 3E**), limited GADD34 upregulation (**Figure 3H**), and decreased CReP expression (**Figure 6E**). We hypothesize that SARS-CoV-2 may benefit from eIF2α phosphorylation, and thus may both induce phosphorylation and limit dephosphorylation to maximize cellular translational shutoff. How SARS-CoV-2 can escape the negative effects of p-eIF2α while other betacoronaviruses cannot remains to be determined. It is possible that SARS-CoV-2 has evolved a way to promote localized dephosphorylation of p-eIF2α around viral mRNAs (57), thus promoting even further skewing of cellular translation towards viral mRNAs. Additionally, nsp1, the viral replicase protein that interacts with host ribosomes and promotes the selective translation of viral mRNAs (51), could play a role. Indeed, a recent study found that SARS-CoV-2 nsp1 binds to the initiation factors EIF1 and EIF1A to enhance the translation of viral transcripts (58). Mechanisms such as this, as well as other undiscovered functions of SARS-CoV-2 replicase and accessory proteins, could help to keep viral translation rates high under conditions of a translationally limited host. Possibly, SARS-CoV-2 may play mediate this process through nsp1, while HCoV-OC43 or MERS-CoV, which also encode nsp1, would not have this capability, a question for future investigation.

It is surprising and unorthodox that CReP, which promotes continuous, low-level dephosphorylation, could compensate for the loss of GADD34 during intense ER stress, such as during coronavirus infection. However, studies that have suggested that CReP has a limited capability to compensate for GADD34 (10, 57, 59) did not include viral infection, which could alter typical function. For instance, during SARS-CoV-2 infection, we observed decreased CReP expression at the mRNA level (**Figure 6E**) and protein level (**Figure 6C**). Additionally, SARS-CoV-2 induced the lowest levels of GADD34 compared to HCoV-OC43 (compare **Figures 3H** and **3I**) and MERS-CoV (compare **Figures 3H** and **3G**). Thus, we conclude that HCoV-OC43 induces both GADD34 and CReP during infection, maximizing eIF2α dephosphorylation to maintain virus protein production. SARS-CoV-2, on the other hand, induces low levels of GADD34 and even decreases CReP levels, thus allowing continued eIF2α phosphorylation throughout infection while somehow not affecting SARS-CoV-2 protein production. MERS-CoV lies somewhere in the middle, relying on eIF2α dephosphorylation, but not to the same extent as HCoV-OC43. Targeting both GADD34 and CReP with salubrinal (32) may serve as an effective therapeutic against MERS-CoV and especially HCoV-OC43.

It remains unclear exactly how HCoV-OC43 and SARS-CoV-2 may be differentially regulating CReP expression during infection. Previous studies have reported CReP can be negatively regulated by the IRE1 pathway of the unfolded protein response via regulated IRE1-dependent decay (RIDD), which degrades CReP mRNA (60). However, we have previously reported that HCoV-OC43 strongly activates IRE1 during infection, while SARS-CoV-2 inhibits the activation of the IRE1 RNase domain (34). This would be expected to produce the opposite regulation of CReP to that we observed during HCoV-OC43 and SARS-CoV-2 infection if RIDD were indeed involved (**Figure 6E**). CReP has also been found to be negatively regulated by mir-98-5p (61, 62), which could be investigated in future studies as a possible mechanism for SARS-CoV-2 reducing CReP expression during infection. While the exact mechanism of CReP upregulation is unclear, it has been reported that CReP mRNA levels can increase to compensate for GADD34 loss under stress conditions, indicating that CreP expression might not always be constitutive (10). We hypothesize that HCoV-OC43 induces such extreme levels of ER stress that this triggers the upregulation of not only GADD34 but also CReP as well. However, further studies will be necessary to unravel this connection.

While our findings regarding GADD34 and CReP during betacoronavirus infection are novel, other groups have reported on the role of PERK during MERS-CoV infection. These publications have concluded that MERS-CoV activates PERK during infection, leading to apoptosis through CHOP upregulation. Interestingly, they found that apoptosis mediated by PERK is beneficial to MERS-CoV replication, but not to SARS-CoV-2 (40), and PERK inhibitors are potentially antiviral to MERS-CoV (39). This demonstrates that MERS-CoV must balance the negative impacts of PERK activation – eIF2α phosphorylation – to exploit this pathway, further supporting the potential efficacy of host-directed therapeutics. This further demonstrates that CoV interactions with the UPR are exceedingly complex and that there is much more to be explored regarding the PERK pathway and its intricate connections to translation, ER health, and cell fate.

Based on our findings, we propose eIF2α dephosphorylation as a potential host-directed therapeutic target during embeco- or merbecovirus infection. Salubrinal treatment led to reductions in MERS-CoV and HCoV-OC43 replication, while CReP depletion confirmed that this protein is necessary for optimal HCoV-OC43 replication and eIF2α dephosphorylation. Interestingly, HCoV-OC43 seems to require inhibition of both GADD34 and CReP to maximally reduce viral titers. Deletion of both GADD34 and CReP has been reported to be toxic to cells. In the case of GADD34 or CReP loss alone, the other can compensate and enable cell survival under conditions of stress (10). Deletion of both prevents all eIF2α dephosphorylation and thus brings the ternary complex concentration to toxically low levels (45, 59), which likely explains why we could not produce a double knockout cell line and limits the usefulness of long-term salubrinal treatment. Thus, single-target inhibitors such as Sephin1 (45) for GADD34 or Raphin1 (59) for CReP would be necessary for *in vivo* treatments, while limited doses or treatment courses of drugs such as salubrinal could be considered. Targeting ER stress has been proposed as a therapeutic strategy for coronaviruses before, with PERK inhibitors (39, 63) (as discussed above), Tg (38), and TM analogs (64) being reported to be effective at combating coronavirus replication in cells. However, stress-inducing drugs are likely to have systemic toxicity (42) that, in cases of severe CoV infection, could harm already stressed organs. As viruses are much more sensitive to translational perturbations than their hosts (65-67), it is possible that rapid treatment with GADD34 inhibitors could deliver a host-directed antiviral effect that primarily targets infected cells. However, our understanding of the interactions of coronaviruses with translation, eIF2α phosphorylation, and host cell stress responses is still in a very early stage, and much more work remains to be done.

## MATERIALS AND METHODS

### Cell Lines

Human A549 cells (ATCC CCL-185) and its derivatives were cultured in RPMI 1640 (Gibco catalog no. 11875) supplemented with 10% fetal bovine serum (FBS), 100 U/mL penicillin, and 100 mg/mL streptomycin (Gibco catalog no. 15140). African green monkey kidney Vero cells (E6) (ATCC CRL-1586) and VeroCCL81 cells (ATCC CCL-81) were cultured in Dulbecco’s modified Eagle’s medium (DMEM; Gibco catalog no. 11965) supplemented with 10% FBS, 100 U/mL of penicillin, 100 mg/mL streptomycin, 50 mg/mL gentamicin (Gibco catalog no. 15750), 1 mM sodium pyruvate (Gibco catalog no. 11360), and 10 mM HEPES (Gibco catalog no. 15630). Human Calu-3 cells (ATCC HTB-55) were cultured in DMEM supplemented with 20% FBS without antibiotics. A549^DPP4^(19) and A549^ACE2^(18) cells were generated as described previously. CRISPR-Cas9 knockout cell lines were generated using lentiviruses. Lentivirus stocks were generated by using lentiCRISPR v2 (Addgene #42230) with single guide RNA (sgRNA) targeting GADD34 (AAGGTTCTGATAAGAACCCA) or scrambled sequence (TTCTCCGAACGTGTCACGT).

### Viruses

SARS-CoV-2 (USA-WA1/2020) was obtained from BEI Resources, NIAID, NIH and propagated in VeroE6-TMPRSS2 cells. The genomic RNA was sequenced and found to be identical to that of GenBank version no. MN985325.1. Recombinant MERS-CoV was described previously(20) and propagated in VeroCCL81 cells. SARS-CoV-2 and MERS-CoV infections were performed in a biosafety level 3 (BSL-3) laboratory under BSL-3 conditions, using appropriate and approved personal protective equipment and protocols. HCoV-OC43 was obtained from ATCC (VR-1558) and grown and titrated on VeroE6 cells at 33°C.

### Viral growth kinetics and titration

SARS-CoV-2, MERS-CoV, and HCoV-OC43 infections and plaque assays were performed as previously described(34). In brief, A549 cells were seeded at 3 x 10^5^ cells per well in a 12-well plate for infections. Calu-3 cells were seeded similarly onto rat tail collagen type I-coated plates (Corning no. 356500). Cells were washed once with phosphate-buffered saline (PBS) before being infected with virus diluted in serum-free medium—RPMI for A549 cells or DMEM for Calu-3 cells. Virus was absorbed for 1h at 37°C before the cells were washed 3 times with PBS and the medium was replaced with 2% FBS RPMI (A549 cells) or 4% FBS DMEM (Calu-3 cells). At the indicated time points, 200 mL of medium was collected to quantify released virus by plaque assay and stored at -80°C. For HCoV-OC43 infections, similar infection conditions and media were used; however, virus was absorbed, and the infections were incubated at 33°C rather than 37°C.

Plaque assays were performed using VeroE6 cells for SARS-CoV-2 and HCoV-OC43 and VeroCCL81 cells for MERS-CoV. SARS-CoV-2 and MERS-CoV plaque assays were performed in 12-well plates at 37°C. HCoV-OC43 plaque assays were performed in 6-well plates at 33°C. In all cases, virus was absorbed onto cells for 1h at the indicated temperatures before overlay was added. A liquid overlay was used (DMEM with 2% FBS, 1x sodium pyruvate, and 0.1% agarose). Cell monolayers were fixed with 4% paraformaldehyde and stained with 1% crystal violet after the following incubation times: SARS-CoV-2 and MERS-CoV, 3 days; HCoV-OC43, 5 days. All plaque assays were performed in biological triplicate and technical duplicate.

### Pharmacologic agents

Tunicamycin (Sigma-Aldrich catalog no. T7765) and thapsigargin (Sigma-Aldrich catalog no. T9033) were purchased at >98% purity. For use in tissue culture, tunicamycin and thapsigargin stock solutions were prepared by dissolving in sterile dimethyl sulfoxide (DMSO). Salubrinal (catalog no. HY-15486) and Sal003 (catalog no. HY-15969) were purchased from MedChemExpress, and stock solutions prepared in DMSO. Both compounds were diluted to the desired concentration in media and filtered sterilized before use in cell culture.

### Immunoblotting

Cells were washed once with ice-cold PBS, and lysates were harvested at the indicated times post infection with lysis buffer (1% NP-40, 2 mM EDTA, 10% glycerol, 150 mM NaCl, 50 mM Tris HCl, pH 8.0) supplemented with protease inhibitors (Roche complete mini-EDTA-free protease inhibitor) and phosphatase inhibitors (Roche PhosStop easy pack). After 5 min, lysates were collected and mixed 3:1 with 4x Laemmli sample buffer (Bio-Rad 1610747). Samples were heated at 95°C for 10 min and then separated on SDS-PAGE and transferred to polyvinylidene difluoride (PVDF) membranes. Blots were blocked with 5% nonfat milk and probed with antibodies (Table 1) diluted in the same blocking buffer. Primary antibodies were incubated overnight at 4°C or for 1h at room temperature. All secondary antibody incubation steps were done for 1h at room temperature. Blots were visualized using Thermo Scientific SuperSignal chemiluminescent substrates (catalog no. 34095 or 34080).

### PhosTag Immunoblotting

7% acrylamide gels were poured containing 50μM Phosbind acrylamide (ApexBio F4002) and 100μM Mn^2+^. Equal volumes of samples were loaded into each well and run alongside an EDTA free protein marker (ApexBio F4005) at 100V for approximately 3 hours. Gels were washed 3 times in transfer buffer with 10% methanol and 10mM EDTA for 20 minutes each. Three more washes of 10 minutes each with transfer buffer not containing EDTA were then performed. Transfers were performed as above with a 10% methanol transfer buffer. Proteins imaged as above using the PERK antibody indicated in Table 4.1.

### RNA sequencing

Raw FastQ files were obtained from Gene Expression Omnibus database (GSE193169). Read quality was assessed using FastQC v0.11.2(68). Raw sequencing reads from each sample were quality and adapter trimmed using BBDuk 38.73(69). The reads were mapped to the human genome (hg38 with Ensembl v98 annotation) using Salmon v0.13.1(70). Differential expression between mock, 24 hpi, and 36 hpi experimental conditions were analyzed using the raw gene counts files by DESeq2 v1.22.1(71). Volcano plots were generated using EnhancedVolcano v1.14.0(72).

### Gene set enrichment analyses

Gene set enrichment analysis (GSEA) was used to identify the upregulation of cellular pathways and responses. fgsea v1.22.0(73) was used to perform specific gene set enrichment analyses and calculate normalized enrichment score (NES) and p-adjusted values on each dataset using DESeq2 stat values. Specific enrichment plots for the Reactome Unfolded Protein Response gene set (stable identifier R-HSA-381119) were generated using fgsea.

### Statistical Analysis

All statistical analyses and plotting of data were performed using GraphPad Prism software. RT-qPCR data were analyzed by one-way ANOVA. Plaque assay data were analyzed by two-way analysis of variance (ANOVA) with multiple-comparison correction. Displayed significance is determined by the P value; *, P < 0.05; **, P < 0.01; ***, P < 0.001; ****, P < 0.0001; ns, not significant.

Quantitative PCR (RT-qPCR). Cells were lysed with RLT Plus buffer, and total RNA was extracted using the RNeasy Plus minikit (Qiagen). RNA was reverse transcribed into cDNA with a high-capacity cDNA reverse transcriptase kit (Applied Biosystems 4387406). cDNA samples were diluted in molecular biology-grade water and amplified using specific RT-qPCR primers (see Table 2). RT-qPCR experiments were performed on a Roche LightCycler 96 instrument. SYBR green supermix was from Bio-Rad. Host gene expression displayed as the fold change over mock-infected samples was generated by first normalizing cycle threshold (C_T_) values to 18S rRNA to generate ΔC_T_ values (ΔC_T_ = C_T_ gene of interest - C_T_ 18S rRNA). Next, Δ(ΔC_T_) values were determined by subtracting the mock-infected ΔC_T_ values from the virus-infected samples. Technical triplicates were averaged and means displayed using the equation 2^-Δ(ΔCt)^. Primer sequences are listed in Table 2.

## ACKNOWLEDGEMENTS

We thank Dr. Alejandra Fausto for her help with techniques for culturing HCoV-OC43, Dr. Nicole Bracci for help with drug treatments, scientific discussions and editing of the manuscript, and Dr. Clayton Otter for helpful discussions. We would also like to thank Dr. Michael Beers and Dr. Aditi Murthy for their expertise on the UPR pathways and for reviewing and discussing experiments and data. This work was supported by a grant from the NIH, R01-AI140442. D.M.R. was supported in part by T32 AI055400.

---

**Table S1.** Antibodies.

**Figure 8**
| **Table S1 Antibodies** |  |  |  |  |
| --- | --- | --- | --- | --- |
| **Primary Antibody** | **Antibody Species** | **Blocking Buffer** | **Dilution** | **Catalog Number** |
| PERK | rabbit | 5% milk | 1:1000 | Cell Signaling Technology 3192S |
| pPKR (phospho-T446) [E120] | rabbit | 5% milk | 1:1000 | Abcam 32036 |
| PKR (D7F7) | rabbit | 5% milk | 1:1000 | Cell Signaling Technology 12297S |
| p-eif2α (S51) | rabbit | 5% milk | 1:1000 | Cell Signaling Technology 9721S |
| eif2α | rabbit | 5% milk | 1:1000 | Cell Signaling Technology 9722S |
| GADD34 | rabbit | 5% milk | 1:600 | 10449-1-AP (Protein Tech) |
| CReP | rabbit | 5% milk | 1:1000 | 14634-1-AP (Protein Tech) |
| GAPDH (14C10) | rabbit | 5% milk | 1:2000 | Cell Signaling Technology 2118S |
| SARS-CoV-2 N | rabbit | 5% milk | 1:2000 | GTX135357 (Gentex) |
| MERS-CoV N | mouse | 5% milk | 1:2000 | 40068-MM10 (Sino Biological) |
| HCoV-OC43 N | rabbit | 5% milk | 1:2000 | 40643-T62 (Sino Biological) |
| **Secondary Antibody** |  |  |  |  |
| goat anti-rabbit IgG | HRP linked | same as primary | 1:3000 | Cell Signaling Technology 7074S |
| goat anti-mouse IgG | HRP linked | same as primary | 1:3000 | Cell Signaling Technology 7076S |

**Table S2.**
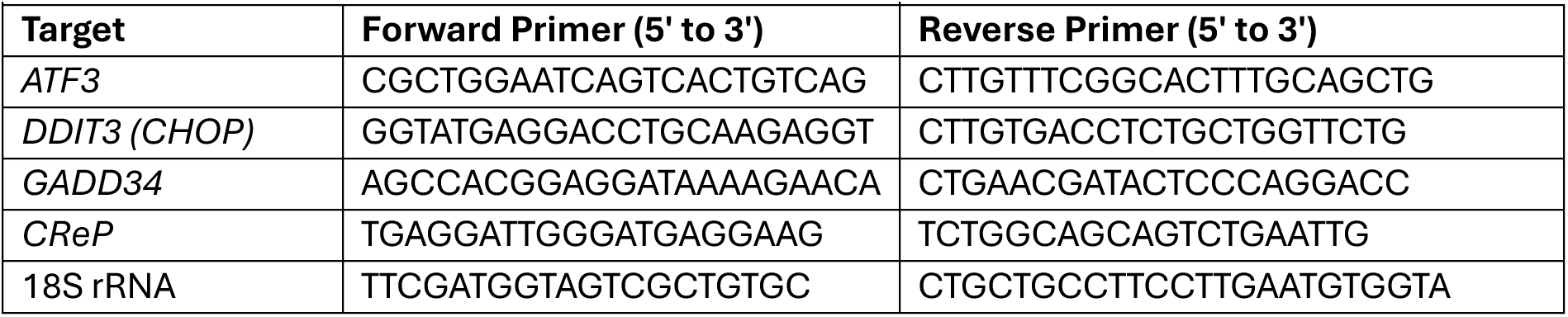
Oligonucleotide primers.

**Figure S1:**
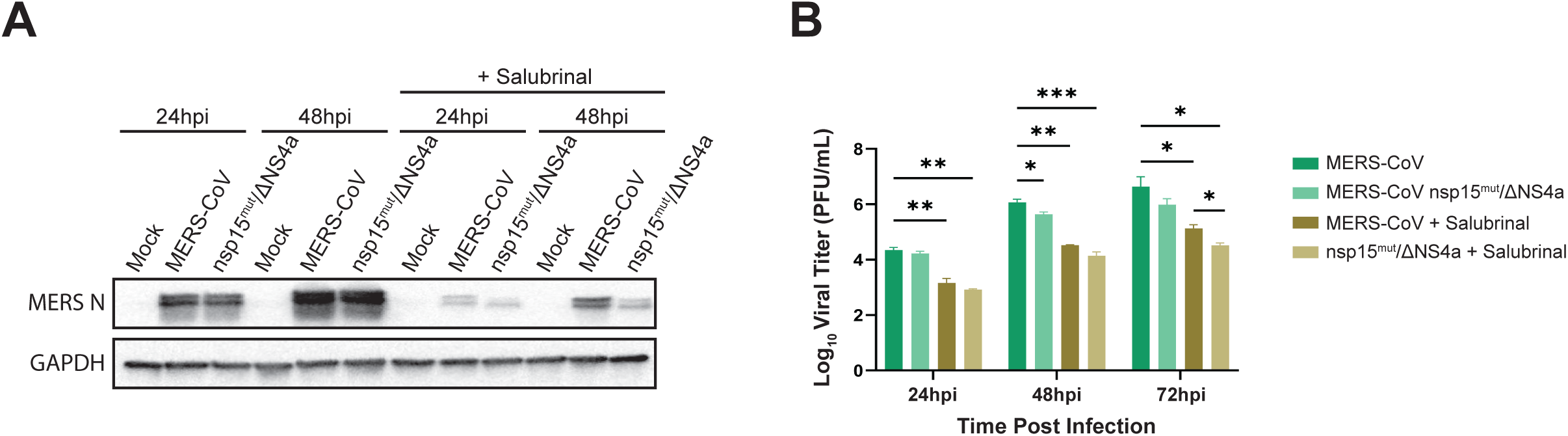
Salubrinal treatment reduces WT MERS-CoV and a MERS-CoV mutant virus replication and translation. A549^DPP4^ cells were infected with MERS-CoV WT or MERS-CoV nsp15^mut^/ΔNS4a at MOI = 0.1. Immediately following infection, cells were left untreated or treated with 20μM salubrinal for the course of the infection. (A) Infectious virus was quantified by plaque assay of supernatants collected from infected cells. (B) Immunoblotting was performed for the indicated proteins, using viral N as a readout for viral translation. Statistics by 2-way ANOVA. * = p < 0.05; ** = p < 0.01; *** p < 0.001; **** = p < 0.0001.

**Figure S2:**
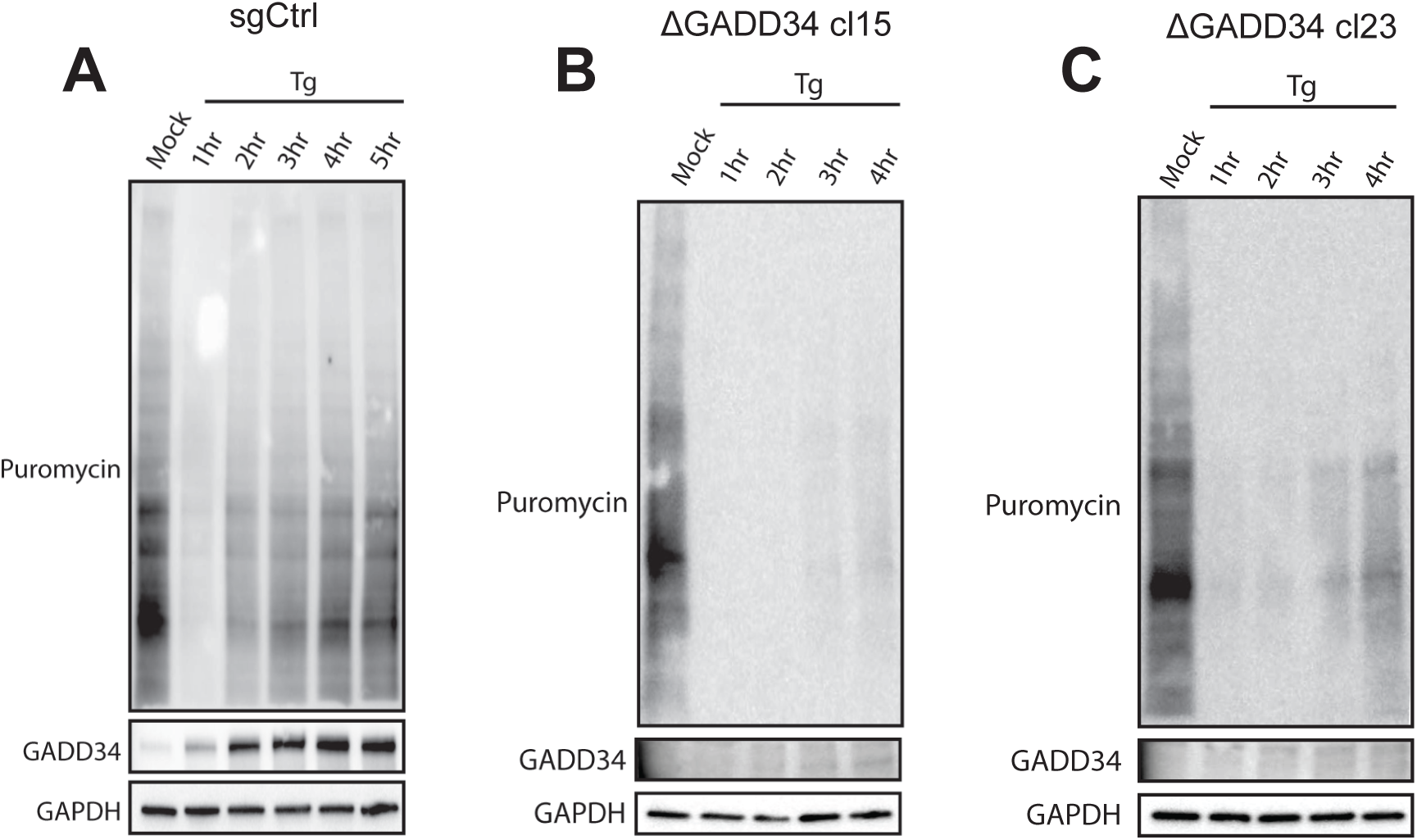
GADD34 knockout reduces translational recovery. A549^ACE2^ single-cell clones with GADD34 knocked out (B and C) or a scramble guide RNA (sgCtrl, A) were treated with 1μM thapsigargin (Tg) or mock treated. At the indicated times, 10μg/mL puromycin was added to the media for 10 minutes before cells were lysed and whole-cell lysates collected. Western immunoblots were performed for the indicated proteins or for puromycin. Cells transduced with scrambled sgRNA (sgCtrl) show rapid GADD34 accumulation and a resumption of translation after 2 hours of Tg treatment. ΔGADD34 cells fail to produce GADD34 protein or restart translation.

## Notes

https://www.ncbi.nlm.nih.gov/geo/query/acc.cgi?acc=GSE193169

### Competing Interest Statement

The authors have declared no competing interest.

